# HLA upregulation during dengue virus infection suppresses the natural killer cell response

**DOI:** 10.1101/698803

**Authors:** Julia L. McKechnie, Davis Beltran, Arcelys Pitti, Lisseth Saenz, Ana B. Araúz, Rosemary Vergara, Eva Harris, Lewis L. Lanier, Catherine A. Blish, Sandra López-Vergès

## Abstract

Dengue virus (DENV) is the most prevalent mosquito-borne virus in the world and a major cause of morbidity in the tropics and subtropics. Upregulation of HLA class I molecules has long been considered a feature of DENV infection, yet this has not been evaluated in the setting of natural infection. Natural killer (NK) cells, an innate immune cell subset critical for mounting an early response to viral infection, are inhibited by self HLA class I, suggesting that upregulation of HLA class I during DENV infection could dampen the NK cell response. Here we addressed whether upregulation of HLA class I molecules occurs during *in vivo* DENV infection and, if so, whether this suppresses the NK cell response. We found that HLA class I expression was indeed upregulated during acute DENV infection across multiple cell lineages *in vivo*. To better understand the role of HLA class I upregulation, we infected primary human monocytes, a major target of DENV infection, *in vitro*. Upregulation of total HLA class I is dependent on active viral replication and is mediated in part by cytokines and other soluble factors induced by infection, while upregulation of HLA-E occurs in the presence of replication-incompetent virus. Importantly, blocking DENV-infected monocytes with a pan-HLA class I Fab nearly doubles the frequency of degranulating NK cells, while blocking HLA-E does not significantly improve the NK cell response. These findings demonstrate that upregulation of HLA class I during DENV infection suppresses the NK cell response, potentially contributing to disease pathogenesis.

## 1 Introduction

Dengue virus (DENV) is a positive-strand RNA virus of which there are four serotypes (DENV-1 to DENV-4). The virus is transmitted between humans by its vector, *Aedes* mosquitoes. Each year, an estimated 390 million people are infected with DENV (Bhatt et al., 2013). While most DENV infections are not life-threatening, severe infections can result in hemorrhage, plasma leakage, shock, organ failure, and death (Kyle and Harris, 2008). The incidence of dengue is rapidly rising (Organization and Others, 2012), increasing the need for a better understanding of how the human immune system responds to DENV infection. There is significant interest in elucidating the role of natural killer (NK) cells during DENV infection. NK cells are innate lymphoid cells that play a key role during the early stages of viral infection. Previous studies have shown that NK cells are activated *in vivo* during DENV infection (Azeredo, 2006; Petitdemange et al., 2016) and that activated NK cells may be an indicator of a positive prognosis (Azeredo, 2006). NK cell activation in response to virally infected cells is dependent on the balance of activating and inhibitory signals from numerous germline-encoded receptors. One such activating receptor, FcRγIIIa (CD16a), mediates antibody-dependent cell cytotoxicity (ADCC), a key bridge between the adaptive and innate immune systems in which antibodies bound to infected cells target them for NK cell killing (Laoprasopwattana et al., 2007; Sun et al., 2017, 2019). NK cells can also kill DENV-infected cells in the absence of ADCC (Costa et al., 2017). Several NK cell receptors, namely DNAM-1, NKG2D, and NKp44 have been implicated in this direct recognition of DENV-infected cells (Beltrán and López-Vergès, 2014; Costa et al., 2017; Mathew, 2018; Petitdemange et al., 2014). However, DENV may also evade the NK cell response, most notably through upregulation of HLA class I (Drews et al., 2018; Glasner et al., 2017; Hershkovitz et al., 2008; Lobigs et al., 1996; Momburg et al., 2001).

HLA class I molecules can bind inhibitory NK cell receptors, mitigating NK cell effector functions against healthy cells. The classical HLA-A, -B, and -C molecules do this by binding to various inhibitory killer-cell immunoglobulin-like receptors (KIRs). The non-classical HLA-E, which presents peptides derived from leader sequences of other HLA molecules, does this by binding to the inhibitory heterodimer CD94/NKG2A (Braud et al., 1998). Viruses can evade NK cell recognition by taking advantage of these inhibitory interactions. *In vitro* studies have shown flaviviruses, including DENV, upregulate total HLA class I as well as HLA-E, leading to inhibition of NK cell activation (Drews et al., 2018; Glasner et al., 2017; Hershkovitz et al., 2008; Lobigs et al., 1996; Momburg et al., 2001). Immune cells, particularly monocytes, are the main targets of DENV infection *in vivo* (Durbin et al., 2008). However, previous studies investigating DENV-mediated HLA class I upregulation and its effect on NK cell activation have used mouse and human cell lines derived from non-immune cells or differentiated primary immune cells (Cheng et al., 2004; Drews et al., 2018; Glasner et al., 2017; Hershkovitz et al., 2008; Libraty et al., 2001; Lobigs et al., 1996; Momburg et al., 2001; Nightingale et al., 2008; Shwetank et al., 2013). This has left a critical gap in our understanding of how undifferentiated primary human immune cell expression of HLA class I is affected by DENV infection, and whether any such changes impact NK cell responses to DENV.

We aimed to determine whether upregulation of class I HLAs, including HLA-E, occurs during *in vivo* DENV infection and, if so, whether this serves to suppress the NK cell response. To address this question, we analyzed peripheral blood mononuclear cell (PBMC) samples from a Panamanian cohort of adult dengue patients and healthy controls for expression of total HLA class I and HLA-E. We then used *in vitro* DENV-infected primary monocytes to determine mediators of HLA class I upregulation. Finally, we co-cultured primary NK cells with autologous, DENV-infected monocytes in the presence of HLA class I blocking Fabs to determine the impact of HLA class I expression on the NK cell response.

## 2 Materials and Methods

### 2.1 DENV patients and ethical statement

Adult DENV patients with less than five days of symptoms consistent with acute DENV infection (fever over 38°C, severe headache, retro-orbital pain, intense myalgia, arthralgia, exanthema, conjunctivitis, diarrhea, chills, nausea, vomiting, abdominal pain, petechiae, and/or bleeding) were recruited at public health institutions (hospitals belonging to the Ministry of Health, the Social Security System in Panama City, Republic of Panama, and suburban areas). Healthy Panamanian control donors volunteered at Gorgas Memorial Institute of Health Studies. All dengue cases were confirmed by qRT-PCR, NS1 antigen, DENV-specific IgM and IgG serological testing. The study protocol was approved by the IRB of Hospital del Niño (CBIHN-M-0634), then confirmed by the committees of ICGES, CSS, Santo Tomas Hospital, and Stanford University. Anonymous healthy adult PBMC samples for *in vitro* studies were collected from leukoreduction system chambers purchased from the Stanford Blood Center.

### 2.2 PBMC sample processing, storage, and thawing

PBMCs were isolated using gradient centrifugation separation by Ficoll-Paque, suspended in freezing media (90% FBS, 10% DMSO), stored at -80°C for 24-72 hours, then transferred to liquid nitrogen. PBMCs were thawed, added to complete media (RPMI-1640, 10% FBS, 1% L-glutamine, 1% penicillin/streptomycin), centrifuged, and counted.

### 2.3 Mass cytometry staining, data acquisition, and analysis

Antibodies for mass cytometry were conjugated using Maxpar^®^ X8 Antibody Labeling Kits (Fluidigm). PBMCs were stained with 25 µM cisplatin (Enzo Life Sciences), live cell palladium barcoded for 30 minutes at 4°C (Mei et al., 2015), pooled, and stained with surface antibodies for 30 minutes at 4°C. Cells were fixed with 2% paraformaldehyde in PBS and permeabilized (eBioscience Permeabilization Buffer) prior to intracellular staining for 45 minutes at 4°C. Finally, the cells were incubated at 4°C in iridium-191/193 intercalator (DVS Sciences) for up to a week, washed once with CyPBS (10X Rockland PBS diluted to 1X in MilliQ water), washed three times with MilliQ water, and diluted with EQ Four Element Calibration Beads before being run on a Helios mass cytometer (Fluidigm). Raw FCS files were normalized using the Normalizer multivariate curve resolution. Normalized files were then de-barcoded using the ParkerICI Premessa de-barcoder. FlowJo^®^ 10.2 was used to gate on live cells. viSNE analysis was performed in Cytobank.

### 2.4 Human sHLA-E ELISA testing

Human sera from 6 DENV confirmed patients and 31 healthy donors were diluted 1:10 and assayed with the Human MHCE/HLA-E ELISA Kit (Biomatik) per manufacturer’s instructions. Optical densities were used to calculate concentration (ng/mL) with a 4 parametric logistic regression analysis using GraphPad Prism 7.

### 2.5 Monocyte and NK cell preparation

Monocytes were isolated from PBMCs by negative selection using a human Pan Monocyte Isolation Kit (Miltenyi). Autologous NK cells were isolated by negative selection using a human NK Cell Isolation Kit (Miltenyi) and cultured in complete RPMI-1640 media with 300 IU/mL of IL-2 (R&D Systems) for 22 hours.

### 2.6 DENV infection of primary monocytes and analysis of HLA expression

*Aedes albopictus* C6/36 cells were infected with DENV-2 laboratory strain 429557 (NR-12216). Supernatants were harvested on day 7 or 8 post-infection, filtered, and ultracentrifuged on a D-sorbitol cushion at 59,439 RCF at 4°C for 3 hours. Virus was titrated using a Vero cell focus-forming assay (Bayless et al., 2016). Concentrated virus was stored at -80°C. Virus was UV-inactivated at 500 μJ x 100 on ice in flat-bottom 96-well plates using a Stratagene UV Stratalinker. Virus inactivation was verified by focus-forming assays. All experiments were repeated with multiple virus batches. Monocytes were mock-infected, exposed to UV-inactivated DENV-2, or infected with active DENV-2 at a multiplicity of infection (MOI) of 2 for 2 hours in infection media (RPMI-1640 media with 2% FBS, 1% penicillin/streptomycin, 1% L-glutamine, and 20 mM HEPES). After 2 hours, cells were washed, resuspended in 24 hour infection media (infection media without HEPES), and incubated at 37°C, 5% CO_2_ for 22 or 46 hours. Cells were stained with FITC-conjugated anti-CD3 (UCHT1, BioLegend), FITC-conjugated anti-CD7 (CD7-6B7, BioLegend), PE-Cy7-conjugated anti-HLA-E (3D12, BioLegend), PE-conjugated anti-pan HLA class I (W6/32, BioLegend), flavivirus group antigen (4G2, Novus Biologicals) conjugated to Alexa Fluor™ 647 using Alexa Fluor 647 Antibody Labeling Kit (Life Technologies), and LIVE/DEAD™ Fixable Yellow Dead Cell Stain Kit (Life Technologies) before analysis on a MACSQuant Analyzer and FlowJo^®^ 10.2.

### 2.7 Supernatant swap assay

Conditioned supernatants from aforementioned DENV-infected primary monocyte cultures were UV-inactivated as previously described. They were then used to culture primary monocytes from the same donors from which the supernatants were collected. After a 24-hour incubation, monocytes were stained with APC-conjugated anti-CD3, APC-conjugated anti-CD7, PE-conjugated anti-pan HLA class I, and Zombie Aqua Fixable Viability dye (BioLegend), then analyzed using a Cytek^™^ Aurora analyzer and FlowJo^®^ 10.2.

### 2.8 Quantification of cytokine production by Luminex

The concentrations of cytokines in conditioned supernatants from the aforementioned DENV-infected primary monocyte cultures were assessed in duplicate using a multiplex cytokine assay by Luminex per the manufacturer’s instructions.

### 2.9 NK cell degranulation assay

Monocytes were infected with DENV-2 at an MOI of 2 and incubated for 24 hours. After incubation, monocytes were left unblocked, blocked with an anti-pan human HLA class I Fab (generated from DX17, BD Bioscience), with an anti-human HLA-E Fab (generated from 3D12, BioLegend), or with isotype-matched control mouse IgG1 Fab (generated from MG1-45, BioLegend) all at 7.3 μg/mL for 30 minutes before adding autologous, IL-2-activated NK cells at a 1:5 effector to target (E:T) ratio. Fabs were produced using mouse IgG1 Fab F(ab)2 Kits (Thermo Scientific) and verified by gel electrophoresis and Coomassie Blue staining. During the 4-hour co-culture, cells were incubated with brefeldin A (eBioscience), monensin (eBioscience), and APC-H7-conjugated anti-CD107a (H4A3, BD Bioscience) per manufacturer’s instructions. Cells were stained with PerCP-Cy5.5-conjugated anti-CD3, FITC-conjugated anti-CD7, Alexa Fluor 700-conjugated anti-CD16 (3G8, BioLegend), PE-Cy7-conjugated anti-CD56 (HCD56, BioLegend), and Zombie Aqua Fixable Viability dye, then analyzed using a Cytek™ Aurora analyzer and FlowJo^®^ 10.2.

### 2.10 Statistical Analysis

A Friedman test, followed by Dunn’s multiple comparisons test was used to determine significant differences between paired data with three conditions. A paired Wilcoxon signed-rank test was used to determine significant differences between DENV- and DENV+ cells. A Friedman test with FDR correction followed by a one-tailed Wilcoxon matched-pairs signed-rank test with a holm correction was used to analyze the Luminex data. Fab blocking data was analyzed using a Friedman test followed by paired Wilcoxon signed-rank tests. All statistical analysis was done using GraphPad Prism 8, R version 3.4.2, R version 3.6.0, and the compare_means function in the open source ggpubr R package.

## 3 Results

We evaluated HLA class I expression on PBMCs from a Panamanian cohort of 8 qRT-PCR confirmed, DENV-2-infected adults within 5 days of symptom onset and 31 healthy Panamanian adult controls **(Supplementary Table 1)**. The expression patterns of HLA class I across cell subsets were visualized with viSNE. This algorithm separated immune cell subsets into clusters based on expression of key lineage markers; manual gating confirmed cluster identity **(Figure 1A and Supplementary Figure 1A)**. Analysis based on protein expression revealed marked upregulation of total HLA class I and HLA-E in DENV-infected adults compared to healthy controls across multiple immune cell subsets **(Figure 1B and Supplementary Figure 1B)**. As HLA-E can also be shed as soluble HLA-E (sHLA-E), which has been implicated as a potential viral mechanism of NK cell evasion (Shwetank et al., 2013, 2014), we performed an sHLA-E ELISA. There was no significant difference in the concentration of sHLA-E between the DENV-infected adults and healthy controls **(Supplementary Figure 2)**. Together, these findings indicate that upregulation of HLA class I occurs on the cell surface of multiple immune cell subsets during acute *in vivo* DENV infection.

**Figure 1:**
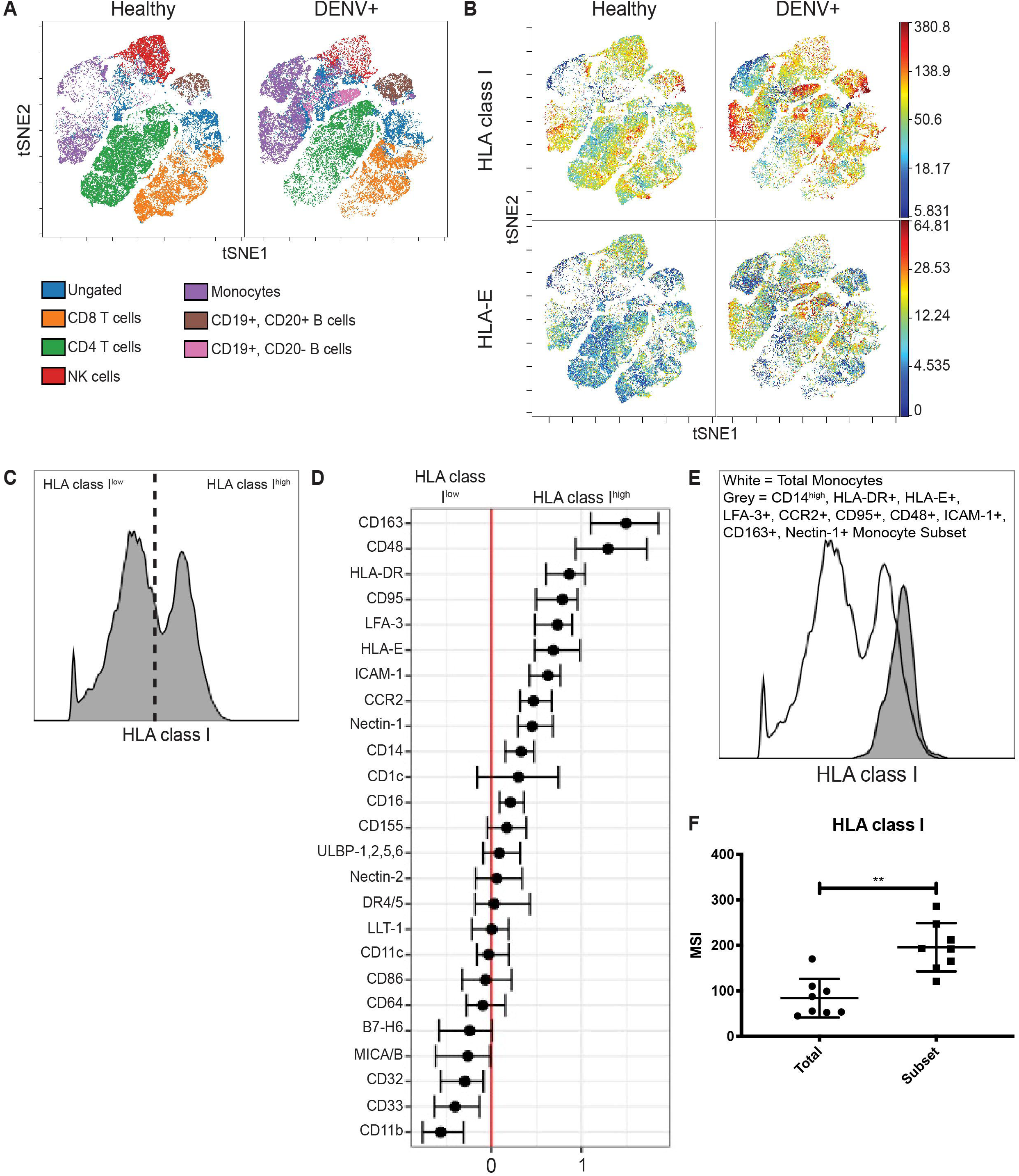
HLA class I upregulation occurs during *in vivo* DENV infection. **(A)** Visualization of immune cell subsets in PBMCs from acute Panamanian DENV patients and healthy Panamanian controls using viSNE. The plots represent pooled data from n = 8 DENV patients and n = 31 controls. To assure equal donor representation, 4375 events were used from each DENV patient and 1129 events were used from each healthy control, resulting in 34,999 pooled events to generate both the DENV+ and healthy control viSNEs. Color key demonstrates major cell populations as determined by gating overlaid upon the viSNE visualization, demonstrating clusters of major cell subsets. **(B)** viSNE visualization of total HLA class I and HLA-E expression in whole PBMCs from DENV patients and healthy controls, generated as in **A. (C)** Representative histogram from a DENV patient illustrating HLA class I^high^ and HLA class I^low^ expressing monocytes gated on for generalized linear mixed model (GLMM) analysis. **(D)** GLMM analysis of markers associated with HLA class I^high^ and HLA class I^low^ expressing monocytes. **(E)** Representative histogram from a DENV patient showing increased HLA class I expression by the monocyte subset gated on using the 10 markers identified in **D** (CD163, CD48, HLA-DR, CD95, LFA-3, HLA-E, ICAM-1, CCR2, Nectin-1, and CD14) compared to total monocytes from the same donor. **(F)** Summary data from all eight DENV patients. Wilcoxon signed-rank test **P < 0.01.

**Figure 2:**
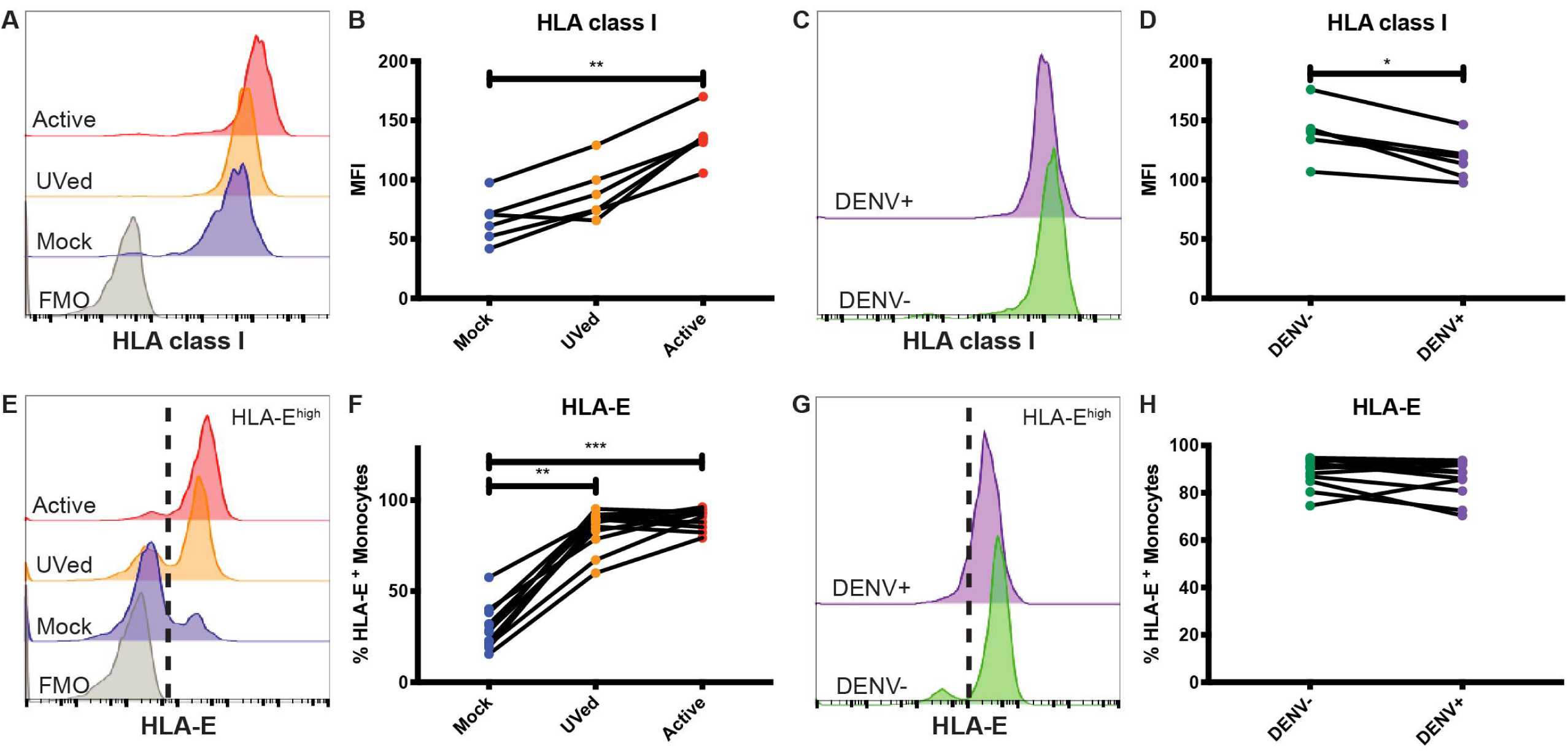
Primary monocytes upregulate HLA class I and HLA-E during *in vitro* DENV infection. Primary monocytes isolated from whole PBMCs from healthy blood bank donors were mock-infected (blue), exposed to UV-inactivated DENV (orange), or infected with active DENV (red) for 48 hours. Representative histograms of HLA class I **(A)** and HLA-E **(E)** expression in total monocytes cultured in the respective conditions. Fluorescence minus one (FMO) shown in grey. HLA class I MFI in total monocytes **(B)** as well as bystander (DENV-) and infected (DENV+) monocytes **(D)**. Representative histograms of HLA class I **(C)** and HLA-E **(G)** expression in bystander monocytes (DENV-, green) and infected monocytes (DENV+, purple). Percentage of HLA-E^high^ total monocytes **(F)** as well as bystander and infected monocytes **(H)**. Two independent experiments measuring HLA class I were performed with 6 donors. The average for each donor is represented in the graphs. n = 12 for HLA-E experiments. Friedman test followed by Dunn’s multiple comparisons test was used to analyze total monocytes. Wilcoxon signed-rank test was used to analyze DENV-vs. DENV+ monocytes. *P < 0.05, **P < 0.01, ***P < 0.0001.

Interestingly, viSNE visualization revealed that upregulation of total HLA class I and HLA-E was not uniform across all monocytes. Instead, there were clear HLA class I^high^ and HLA-E^high^ expressing monocytes. We gated on these cells **(Figure 1C and Supplementary Figure 3A)** and used an unbiased generalized linear mixed model (GLMM) (Seiler et al., 2019) to identify associated markers. The GLMM identified 10 markers (CD14, HLA-DR, HLA-E, LFA-3, CCR2, CD95, CD48, ICAM-1, CD163, and Nectin-1) whose expression was associated with HLA class I^high^ monocytes **(Figure 1D)** and 5 markers (HLA class I, CD11b, ULBP-1,2,5,6, CD163, and MICA/B) whose expression was associated with HLA-E^high^ monocytes **(Supplementary Figure 3B)**. We then verified these markers by comparing the expression level of each marker in HLA^high^ expressing monocytes to the expression level in HLA^low^ expressing monocytes from DENV-infected adults **(Supplementary Figure 3C and Supplementary Figure 4A)**. Finally, we gated down to monocytes expressing all 10 or all 5 markers in DENV-infected adults **(Supplementary Figure 3D and Supplementary Figure 4B)**, and found that the 10 marker subset and the 5 marker subset expressed HLA class I and HLA-E, respectively, at significantly higher levels than total monocytes from the same donor **(Figure 1E, Figure 1F, Supplementary Figure 3E, and Supplementary Figure 3F)** verifying that combinatorial gating using the aforementioned markers accurately identifies HLA class I^high^ and HLA-E^high^ monocytes. Using the same gating scheme, we gated on the 10 and 5 marker monocyte subsets in healthy controls and found that HLA class I expression by the 10 marker subset and HLA-E expression by the 5 marker subset were both 1.6-fold higher in DENV-infected adults compared to healthy controls **(Supplementary Figure 3G and Supplementary Figure 4C)**. These results indicate that upregulation of HLA class I and HLA-E by monocytes during *in vivo* DENV infection only occurs on specific monocyte subsets.

**Figure 3:**
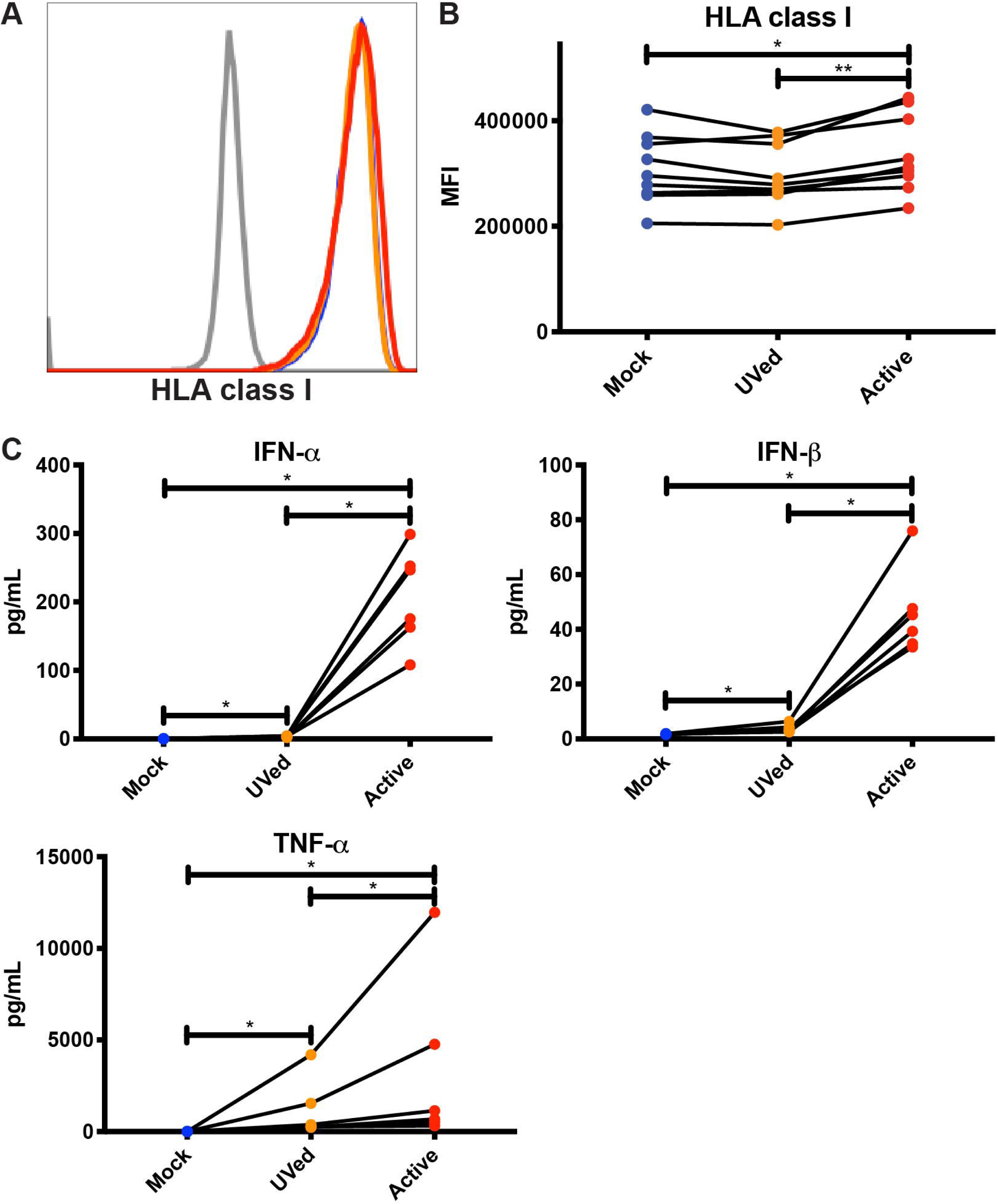
Soluble factors secreted during active DENV infection upregulate HLA class I expression. Conditioned supernatants from experiments shown in Figure 2 were UV-treated and used to culture primary monocytes from healthy blood bank donors (n = 9). After a 24-hour incubation, expression of total HLA class I was analyzed by flow cytometry. Histograms from a single representative donor **(A)** as well as summary data from all 9 donors **(B)** are shown. Friedman test followed by Dunn’s multiple comparisons test, *P < 0.05, **P < 0.01. **(C)** Cytokine concentrations in conditioned supernatants from experiments shown in Figure 2 were analyzed by Luminex. Values shown are the average of two reads for each sample (n = 6). Friedman test with FDR correction followed by a one-tailed Wilcoxon matched-pairs signed-rank test with holm correction, *P < 0.05.

To better understand the effects of HLA class I upregulation, we modeled DENV infection *in vitro* using primary monocytes isolated from healthy donors. Further, to understand how viral replication versus the presence of viral proteins alters HLA class I expression, we compared the effects of ‘active’, replication-competent DENV, with that of UV-inactivated virus incapable of viral replication. At 24 hours post-infection (hpi), HLA class I expression did not significantly differ between active DENV, UV-inactivated DENV, or mock-infected conditions **(Supplementary Figures 5A and 5B)**. By 48 hpi, HLA class I expression was 2.1-fold higher in monocytes infected with active DENV compared to mock **(Figure 2A and 2B)**. Interestingly, in the active virus cultures, the uninfected bystander monocytes had a modest 1.1- and 1.2-fold higher HLA class I expression at 24 **(Supplementary Figures 5C and 5D)** and 48 **(Figure 2C and 2D)** hpi, respectively, than infected monocytes in the same culture. This suggests HLA class I upregulation is primarily restricted to infected cells with a modest impact on uninfected cells, likely due to changes in the cytokine milieu. Alternatively, bystander cells in our assay may have been infected at a level below our limit of detection.

Expression of HLA-E was also altered during DENV infection. At 24 and 48 hpi, exposure to UV-inactivated DENV or infection with active DENV resulted in a majority of monocytes becoming HLA-E^high^ **(Supplementary Figure 5E and Figure 2E)**. Similarly, at 48 hpi, the percentage of HLA-E^high^ monocytes in the UVed and active DENV cultures was 2.8- and 3-fold higher, respectively, compared to mock-infected monocytes **(Figure 2F)**. The increase of HLA-E^high^ monocytes was 3-fold for both virus conditions compared to mock at 24 hpi **(Supplementary Figure 5F)**. At both time points, bystander monocytes in the active DENV cultures had a bimodal distribution of HLA-E^low^ and HLA-E^high^ cells, while infected cells were all HLA-E^high^ **(Figure 2G and Supplementary Figure 5G)**. Infected monocytes displayed a 1.3-fold increase in the percentage of HLA-E^high^ monocytes at 24 hpi compared to bystander monocytes **(Supplementary Figure 5H)**, but at 48 hpi there was no longer a significant difference **(Figure 2H)**. These results suggest that the response to viral proteins, rather than viral replication, is the main driver of HLA-E upregulation.

Considering that bystander monocytes expressed higher levels of HLA class I than DENV-infected monocytes, we wanted to test whether secreted factors, such as cytokines, viral proteins, and other molecules produced during active DENV infection were sufficient to upregulate HLA class I. To this end, we UV-treated conditioned supernatants from previous 48-hour cultures of primary monocytes that were mock-infected, exposed to UV-inactivated DENV, or infected with active DENV. We then isolated uninfected monocytes from the same donors from which the supernatants were collected and cultured them in the conditioned supernatants for 24 hours. Supernatants collected from active, DENV-infected cultures led to a significant, if modest, 1.1-fold increase in HLA class I expression compared to monocytes cultured in supernatants collected from the mock-infected and UV-inactivated DENV cultures **(Figure 3A and 3B)**. This increase in HLA class I expression shows that soluble factors secreted during active DENV infection could be contributing to HLA class I upregulation. In order to investigate the potential role of cytokines in mediating this increase in HLA class I expression, we used Luminex to determine the concentration of cytokines present in the conditioned supernatants. We found that the concentrations of IFN-α, IFN-β, and TNF-α were 759.2-, 26-, and 271.7-fold higher respectively in the active condition compared to mock and 59.9-, 12.3-, and 2.8-fold higher respectively in the active condition compared to the UVed condition **(Figure 3C)**. These findings suggest that upregulation of HLA class I during active DENV infection is largely driven by viral replication, and is likely mediated to some extent by cytokines. It is important to note that viral RNA, proteins, and particles present in the supernatant of active infection cultures could also contribute to HLA class I upregulation.

Given that NK cell activation is dampened by the expression of self-HLA class I on potential target cells, we investigated the impact of DENV-mediated HLA class I upregulation on NK cell degranulation in response to DENV-infected cells. We co-cultured DENV-infected primary monocytes with autologous primary NK cells in the presence of an isotype-matched control Fab, an anti-pan HLA class I blocking Fab, or an anti-HLA-E blocking Fab for 4 hours and measured the percentage of CD107a+ NK cells as a marker of degranulation and killing activity **(Figure 4A and 4B)**. Fabs were used instead of whole IgG to avoid killing via ADCC following binding of the anti-HLA antibodies to the target cells. Blocking HLA class I on mock-infected monocytes led to a 9.7% frequency of CD107a+ NK cells, a 4.5% increase from 5.2% in the unblocked, mock-infected monocyte condition. Because NK cells can become activated when they are unable to bind self-HLA class I, this result demonstrates that the Fabs were effectively blocking their targeted proteins. Blocking HLA class I on DENV-infected monocytes resulted in nearly double the frequency of CD107a+ NK cells compared to DENV-infected monocytes blocked with the isotype-matched control Fab, increasing from 9.9% to 18.8%. Additionally, HLA class I-blocking of DENV-infected monocytes resulted in a 6.7% increase in CD107a+ NK cells compared to blocking HLA-E. Blocking HLA class I on DENV-infected monocytes also nearly doubled the frequency of CD107a+ NK cells compared to blocking HLA class I on mock-infected cells, increasing from 9.7% to 18.8% (P = 0.0312). For all of the blocking conditions, except HLA-E, the DENV co-cultures had a statistically significant increase in CD107a+ NK cells compared to the mock-infected co-cultures. Thus, these data demonstrate that HLA class I upregulation can dampen the NK cell response to DENV-infected cells.

**Figure 4:**
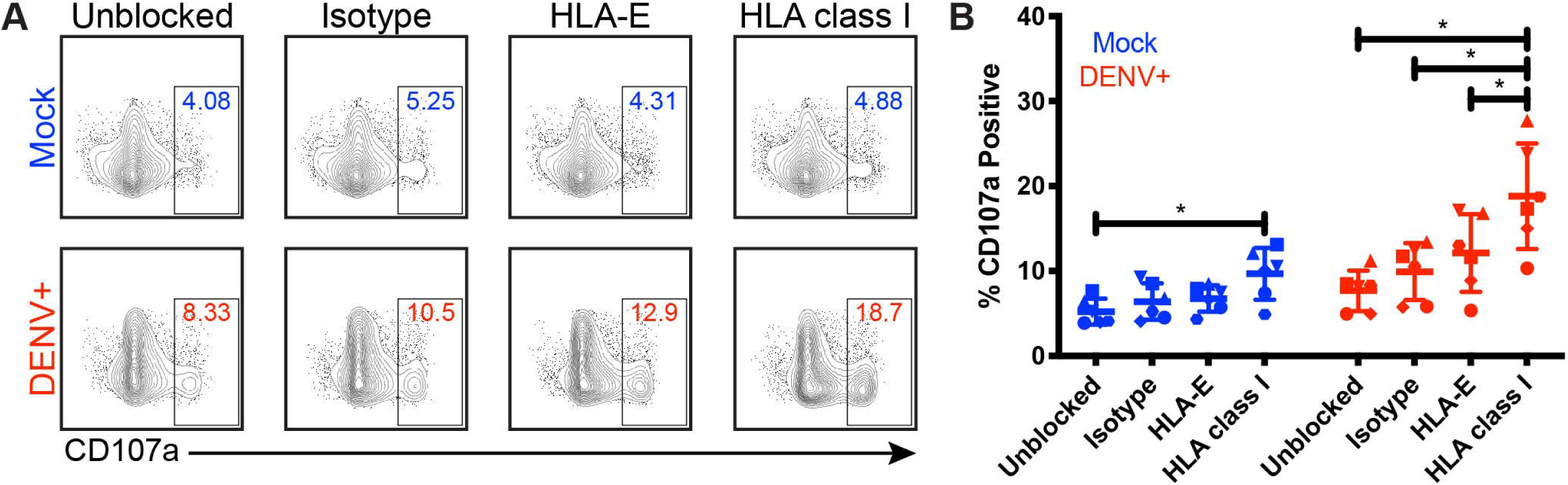
Blocking HLA class I improves NK cell responses to DENV-infected cells. Primary NK cells and monocytes were isolated from whole PBMCs from healthy blood bank donors (n = 6). NK cells were activated for 22 hours with IL-2. Monocytes were mock-infected (blue) or infected with active DENV (red) at an MOI of 2 for 24 hours. Prior to co-culture with autologous NK cells, monocytes were blocked for 30 minutes with an isotype-matched control Fab, an anti-HLA-E blocking Fab, or an anti-pan HLA class I blocking Fab. Monocytes and NK cells were co-cultured for 4 hours before NK cell expression of CD107a was evaluated by flow cytometry. Flow cytometry plots from a single representative donor **(A)** as well as summary data from all 6 donors **(B)** are shown. Friedman test followed by paired Wilcoxon signed-rank tests, *P < 0.05.

## 4 Discussion

Roughly one-third of the world’s population is at risk of acquiring DENV, making it critically important that we elucidate immune factors that contribute both to disease protection and pathogenesis. Mechanisms by which DENV evades the innate immune response by inhibiting the production and signaling of type I IFNs, as well as other aspects of the cellular antiviral response, have been well reported (Green et al., 2014; Morrison et al., 2012). Similarly, pathways that might promote DENV escape from NK cell recognition have been proposed, including a potential role for the upregulation of HLA class I molecules by DENV-infected cells to inhibit the NK cell response by binding inhibitory KIRs or CD94/NKG2A (Beltrán and López-Vergès, 2014; Mathew, 2018; Petitdemange et al., 2014). However, these data have primarily arisen from *in vitro* infection of mouse or human cell lines, rather than more physiologic systems such as natural infection or infection of undifferentiated primary human immune cells. Here, we show natural DENV infection leads to increased expression of HLA class I and HLA-E in adult patients. We also found that soluble factors produced during active infection contributed to HLA class I upregulation. Finally, blocking HLA class I on DENV-infected monocytes enhanced the ability of NK cells to degranulate in response to DENV-infected cells. Together, these findings show HLA class I upregulation during active DENV infection suppresses NK cell degranulation.

HLA class I upregulation during flavivirus infection has been previously described and attributed to various mechanisms such as NFκB activation (Kesson and King, 2001), increased transport of peptides into the endoplasmic reticulum for HLA loading (Momburg et al., 2001), and the presence of IFN-β (Glasner et al., 2017). We show replication-competent virus was required for significant HLA class I upregulation, but that HLA class I expression was highest in uninfected bystander monocytes. Further, supernatants collected from active DENV cultures were able to upregulate HLA class I on uninfected monocytes and contained higher concentrations of IFN-α, IFN-β, and TNF-α compared to supernatants from mock-infected or UV-inactivated DENV cultures indicating HLA class I upregulation is mediated at least in part by soluble factors. These findings pose a potential mechanism for HLA class I upregulation in which soluble factors secreted by DENV-infected cells induce increased HLA class I expression on all cells in an effort to promote cytotoxic T lymphocyte responses. However, the DENV-infected cells express lower levels than the bystander cells because DENV may encode proteins that interfere with processes driving HLA class I upregulation to escape the T cell response (Glasner et al., 2017; Green et al., 2014; Guzman and Harris, 2015; Ye et al., 2013).

Interestingly, upregulation of HLA-E likely involves different mechanisms than upregulation of other HLA class I molecules. We observed a significant increase in HLA-E expression in response to active DENV as well as UV-inactivated DENV. This suggests that direct sensing of viral products by innate immune receptors and the resulting cytokines, rather than pathways induced during viral replication, may be the primary contributors to HLA-E upregulation. However, DENV-infected monocytes expressed the highest levels of HLA-E, implying DENV itself is also mediating HLA-E upregulation. Intriguingly, cytomegalovirus proteins can modulate the surface expression of HLA-E by encoding HLA leader sequence mimics with reduced binding affinity to the CD94/NKG2 receptors (Heatley et al., 2013). Similarly, human immunodeficiency virus-1 has been found to upregulate HLA-E expression, resulting in the presentation of a capsid peptide which prevents HLA-E engagement with CD94/NKG2A (Davis et al., 2016; Nattermann et al., 2005). No such mechanisms for modulating HLA-E expression and its affinity for the CD94/NKG2 receptors have been reported for DENV. The NetMHCpan 4.0 server predicts several DENV-2 peptides with strong and weak binding to HLA-E*01:01, making it possible that DENV peptides modulate NKG2A/C binding. Together, these results suggest HLA-E upregulation is mediated by both virus-dependent and virus-independent mechanisms, and could influence NK cell recognition through NKG2A/C.

Here we extend prior studies demonstrating that HLA class I upregulation during flavivirus infection in cell lines can inhibit NK cell activation (Drews et al., 2018; Glasner et al., 2017; Hershkovitz et al., 2008; Lobigs et al., 1996; Momburg et al., 2001). For the first time, we used patient samples and more physiologically relevant *in vitro* infection and co-culture systems with undifferentiated primary human immune cells. By directly blocking NK cell binding to HLA class I using Fabs, we found HLA class I expression on DENV-infected cells significantly dampens NK cell degranulation. This provides the first direct evidence that upregulation of HLA class I is responsible for inhibition of NK cell responses to flavivirus-infected cells. We did observe some non-specific increase in NK cell degranulation in the presence of the isotype-matched control antibody, but this effect was dwarfed by the increase in NK cell degranulation in response to DENV-infected cells in the presence of HLA class I blocking Fabs. These findings indicate that HLA class I expression dampens the magnitude of the NK cell response.

Previous studies using cytokine-activated endothelial cells showed that blocking surface HLA-E and sHLA-E increased NK cell killing, illustrating the importance of HLA-E expression as a potential NK escape mechanism in some vascular diseases (Coupel et al., 2007). However, similar to Drews *et. al.*, we did not observe a significant modulating effect of HLA-E expression on NK cell activity against DENV-infected cells (Drews et al., 2018). HLA-E binds to both inhibitory NK cell receptor CD94/NKG2A and activating receptor CD94/NKG2C (Braud et al., 1998; Kaiser et al., 2005; Valés-Gómez et al., 1999). Notably, HLA-E binds to CD94/NKG2A with higher affinity (Kaiser et al., 2005; Valés-Gómez et al., 1999). The fact that NK cell binding to HLA-E can result in both activating and inhibitory signaling with a dominant advantage towards inhibitory signaling could explain our results and those of Drews *et al*. HLA-E’s greater affinity towards CD94/NKG2A also suggests that increased HLA-E expression on DENV-infected cells might be part of the viral escape strategy. Specifically blocking NKG2A or NKG2C in NK cell-infected cell co-cultures could identify the role of inhibitory signaling through NKG2A versus activating signaling through NKG2C in DENV recognition.

In contrast to Drews *et al.* and Shwetank *et al.* who observed an increase in sHLA-E in the supernatants of DENV-infected HMEC-1 cells and Japanese encephalitis virus-infected human brain microvascular endothelial cells, respectively, we saw no significant increase in sHLA-E in the serum of DENV patients compared to healthy controls (Drews et al., 2018; Shwetank et al., 2013). This suggests that sHLA-E shedding is not increased at the systemic level during *in vivo* DENV infection and is consequently unlikely to contribute strongly to suppression of the NK cell response.

This study has limitations, the most significant of which is the small sample size of DENV-infected adults in our cohort. Despite the modest numbers, the conclusions we drew from these *in vivo* data were clear and supported by our *in vitro* experiments using primary human immune cells. The second limitation is that we were unable to clearly determine the role of HLA-E on the NK cell response to DENV infection given its binding to both inhibitory CD94/NKG2A and activating CD94/NKG2C receptors.

To our knowledge, ours is the first study showing upregulation of HLA class I molecules in acute dengue patient samples, suggesting different drivers of HLA-E upregulation versus upregulation of other HLA class I proteins during DENV infection, and showing enhanced primary NK cell degranulation upon blocking HLA class I on primary DENV-infected monocytes. Our *in vivo* HLA class I expression data need to be confirmed with additional DENV cohorts at different time points of the disease, spanning all serotypes and degrees of disease severity, and including pediatric patients. This will be vital to determining what temporal, viral, and age-related factors affect HLA class I upregulation, as well as whether there is a correlation between disease severity and HLA class I expression. Future experiments are also required to determine what factors and pathways mediate HLA class I upregulation in monocytes and how HLA class I expression is modulated in other immune cell subsets. Overall, this study furthers our understanding of the impacts of DENV infection on innate immune cells and their intercellular interactions.

## 6 Author Contributions

Conceptualization, JM, DB, CB, SL-V; Methodology, JM, DB, AP, LS, AA, RV; Formal Analysis, JM, DB, CB, SL-V; Investigation, JM, DB, CB, SL-V; Resources, DB, EH, LL, CB, SL-V; Data Curation, JM, DB, CB, SL-V; Writing – Original Draft Preparation, JM, DB, CB, SL-V; Writing – Review & Editing, EH, LL; Supervision, CB, SL-V; Project Administration, CB, SL-V; Funding Acquisition, DB, CB, SL-V.

## 8 Acknowledgments

We thank all health institutions, patients, and their families, Luis Bonilla from the Blood Bank of Santo Tomas Hospital for providing blood components, and the Stanford Human Immune Monitoring Center. We thank Dr. Taia Wang for critical reading of the manuscript and Dr. Anne-Maud Ferreira for statistical analysis of the Luminex data.

## 9 Data Availability Statement

Requests to access the dataset should be directed to Catherine Blish at.

## References

Azeredo, E. L. (2006). NK cells, displaying early activation, cytotoxicity and adhesion molecules, are associated with mild dengue disease. 345–356.

Bayless, N. L., Greenberg, R. S., Swigut, T., Wysocka, J., and Blish, C. A. (2016). Zika Virus Infection Induces Cranial Neural Crest Cells to Produce Cytokines at Levels Detrimental for Neurogenesis. Cell Host Microbe 20, 423–428.

Beltrán, D., and López-Vergès, S. (2014). NK Cells during Dengue Disease and Their Recognition of Dengue Virus-Infected cells. Front. Immunol. 5, 192.

Bhatt, S., Gething, P. W., Brady, O. J., Messina, J. P., Farlow, A. W., Moyes, C. L., et al. (2013). The global distribution and burden of dengue. Nature 496, 504–507.

Braud, V. M., Allan, D. S., O’Callaghan, C. A., Söderström, K., D’Andrea, A., Ogg, G. S., et al. (1998). HLA-E binds to natural killer cell receptors CD94/NKG2A, B and C. Nature 391, 795–799.

Cheng, Y., King, N. J. C., and Kesson, A. M. (2004). Major histocompatibility complex class I (MHC-I) induction by West Nile virus: involvement of 2 signaling pathways in MHC-I up-regulation. J. Infect. Dis. 189, 658–668.

Costa, V. V., Ye, W., Chen, Q., Teixeira, M. M., Preiser, P., Ooi, E. E., et al. (2017). Dengue Virus-Infected Dendritic Cells, but Not Monocytes, Activate Natural Killer Cells through a Contact-Dependent Mechanism Involving Adhesion Molecules. MBio 8. doi:10.1128/mBio.00741-17.

Coupel, S., Moreau, A., Hamidou, M., Horejsi, V., Soulillou, J.-P., and Charreau, B. (2007). Expression and release of soluble HLA-E is an immunoregulatory feature of endothelial cell activation. Blood 109, 2806–2814.

Davis, Z. B., Cogswell, A., Scott, H., Mertsching, A., Boucau, J., Wambua, D., et al. (2016). A Conserved HIV-1-Derived Peptide Presented by HLA-E Renders Infected T-cells Highly Susceptible to Attack by NKG2A/CD94-Bearing Natural Killer Cells. PLoS Pathog. 12, 1–22.

Drews, E., Adam, A., Htoo, P., Townsley, E., and Mathew, A. (2018). Upregulation of HLA-E by dengue and not Zika viruses. Clinical & Translational Immunology 7, e1039.

Durbin, A. P., Vargas, M. J., Wanionek, K., Hammond, S. N., Gordon, A., Rocha, C., et al. (2008). Phenotyping of peripheral blood mononuclear cells during acute dengue illness demonstrates infection and increased activation of monocytes in severe cases compared to classic dengue fever. Virology 376, 429–435.

Glasner, A., Oiknine-Djian, E., Weisblum, Y., Diab, M., Panet, A., Wolf, D. G., et al. (2017). Zika Virus Escapes NK Cell Detection by Upregulating Major Histocompatibility Complex Class I Molecules. J. Virol. 91. doi:10.1128/JVI.00785-17.

Green, A. M., Beatty, P. R., Hadjilaou, A., and Harris, E. (2014). Innate immunity to dengue virus infection and subversion of antiviral responses. J. Mol. Biol. 426, 1148–1160.

Guzman, M. G., and Harris, E. (2015). Dengue. Lancet. doi:10.1016/S0140-6736(14)60572-9.

Heatley, S. L., Pietra, G., Lin, J., Widjaja, J. M. L., Harpur, C. M., Lester, S., et al. (2013). Polymorphism in human cytomegalovirus UL40 impacts on recognition of human leukocyte antigen-E (HLA-E) by natural killer cells. J. Biol. Chem. 288, 8679–8690.

Hershkovitz, O., Zilka, A., Bar-Ilan, A., Abutbul, S., Davidson, A., Mazzon, M., et al. (2008). Dengue Virus Replicon Expressing the Nonstructural Proteins Suffices To Enhance Membrane Expression of HLA Class I and Inhibit Lysis by Human NK Cells. J. Virol. 82, 7666–7676.

Kaiser, B. K., Barahmand-Pour, F., Paulsene, W., Medley, S., Geraghty, D. E., and Strong, R. K. (2005). Interactions between NKG2x immunoreceptors and HLA-E ligands display overlapping affinities and thermodynamics. J. Immunol. 174, 2878–2884.

Kesson, A. M., and King, N. J. C. (2001). Transcriptional regulation of major histocompatibility complex class I by flavivirus West Nile is dependent on NF-kappa B activation. J. Infect. Dis. 184, 947–954.

Kyle, J. L., and Harris, E. (2008). Global spread and persistence of dengue. Annu. Rev. Microbiol. 62, 71–92.

Laoprasopwattana, K., Libraty, D. H., Endy, T. P., Nisalak, A., Chunsuttiwat, S., Ennis, F. A., et al. (2007). Antibody-dependent cellular cytotoxity mediated by plasma obtained before secondary dengue virus infections: Potential involvement in early control of viral replication. J. Infect. Dis. 195, 1108–1116.

Libraty, D. H., Pichyangkul, S., Ajariyakhajorn, C., Endy, T. P., and Ennis, F. A. (2001). Human Dendritic Cells Are Activated by Dengue Virus Infection : Enhancement by Gamma Interferon and Implications for Disease Pathogenesis. J. Virol. 75, 3501–3508.

Lobigs, M., Blanden, R. V., and Müllbacher, A. (1996). Flavivirus-induced up-regulation of MHC class I antigens; implications for the induction of CD8+ T-cell-mediated autoimmunity. Immunol. Rev. 152, 5–19.

Mathew, A. (2018). Defining the role of NK cells during dengue virus infection. Immunology, 0–2.

Mei, H. E., Leipold, M. D., Schulz, A. R., Chester, C., and Maecker, H. T. (2015). Barcoding of live human peripheral blood mononuclear cells for multiplexed mass cytometry. J. Immunol. 194, 2022–2031.

Momburg, F., Müllbacher, A., and Lobigs, M. (2001). Modulation of Transporter Associated with Antigen Processing (TAP) -Mediated Peptide Import into the Endoplasmic Reticulum by Flavivirus Infection Modulation of Transporter Associated with Antigen Processing (TAP) -Mediated Peptide Import into the End. 75, 5663–5671.

Morrison, J., Aguirre, S., and Fernandez-Sesma, A. (2012). Innate immunity evasion by Dengue virus. Viruses 4, 397–413.

Nattermann, J., Nischalke, H. D., Hofmeister, V., Kupfer, B., Ahlenstiel, G., Feldmann, G., et al. (2005). HIV-1 infection leads to increased HLA-E expression resulting in impaired function of natural killer cells. Antivir. Ther. 10, 95–107.

Nightingale, Z. D., Patkar, C., and Rothman, A. L. (2008). Viral replication and paracrine effects result in distinct, functional responses of dendritic cells following infection with dengue 2 virus. J. Leukoc. Biol. 84, 1028–1038.

Organization, W. H., and Others (2012). Global strategy for dengue prevention and control 2012-2020. Available at: https://apps.who.int/iris/bitstream/handle/10665/75303/9789241504034_eng.pdf.

Petitdemange, C., Wauquier, N., Devilliers, H., Yssel, H., Mombo, I., Caron, M., et al. (2016). Longitudinal Analysis of Natural Killer Cells in Dengue Virus-Infected Patients in Comparison to Chikungunya and Chikungunya/Dengue Virus-Infected Patients. PLoS Negl. Trop. Dis. 10, 1–17.

Petitdemange, C., Wauquier, N., Rey, J., Hervier, B., Leroy, E., and Vieillard, V. (2014). Control of acute dengue virus infection by natural killer cells. 5, 1–5.

Seiler, C., Kronstad, L. M., Simpson, L. J., Le Gars, M., Vendrame, E., Blish, C. A., et al. (2019). Uncertainty Quantification in Multivariate Mixed Models for Mass Cytometry Data. arXiv [stat.AP]. Available at: http://arxiv.org/abs/1903.07976.

Shwetank Date, O. S., Carbone, E., and Manjunath, R. (2014). Inhibition of ERK and proliferation in NK cell lines by soluble HLA-E released from Japanese encephalitis virus infected cells. Immunol. Lett. 162, 94–100.

Shwetank Date, O. S., Kim, K. S., and Manjunath, R. (2013). Infection of human endothelial cells by Japanese encephalitis virus: increased expression and release of soluble HLA-E. PLoS One 8, e79197.

Sun, P., Morrison, B. J., Beckett, C. G., Liang, Z., Nagabhushana, N., Li, A., et al. (2017). NK cell degranulation as a marker for measuring antibody-dependent cytotoxicity in neutralizing and non-neutralizing human sera from dengue patients. J. Immunol. Methods 441, 24–30.

Sun, P., Williams, M., Nagabhushana, N., Jani, V., Defang, G., and Morrison, B. J. (2019). NK Cells Activated through Antibody-Dependent Cell Cytotoxicity and Armed with Degranulation/IFN-γ Production Suppress Antibody-dependent Enhancement of Dengue Viral Infection. Sci. Rep. 9, 1109.

Valés-Gómez, M., Reyburn, H. T., Erskine, R. A., López-Botet, M., and Strominger, J. L. (1999). Kinetics and peptide dependency of the binding of the inhibitory NK receptor CD94/NKG2-A and the activating receptor CD94/NKG2-C to HLA-E. EMBO J. 18, 4250–4260.

Ye, J., Zhu, B., Fu, Z. F., Chen, H., and Cao, S. (2013). Immune evasion strategies of flaviviruses. Vaccine 31, 461–471.

