## Supplementary figures and images for "HLA upregulation during dengue virus infection suppresses the natural killer cell response"

### Supplementary Figure 1

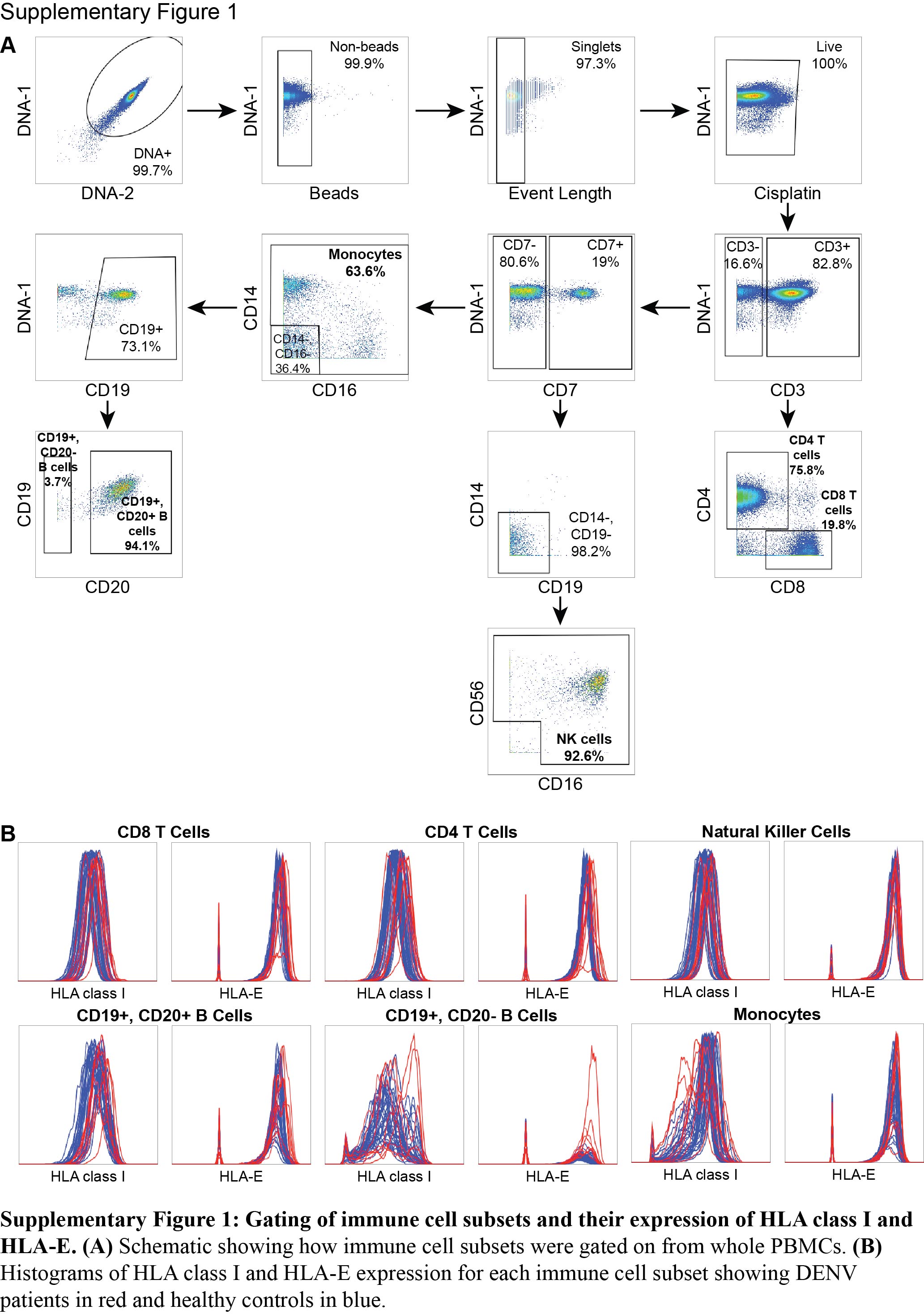

### Supplementary Figure 2

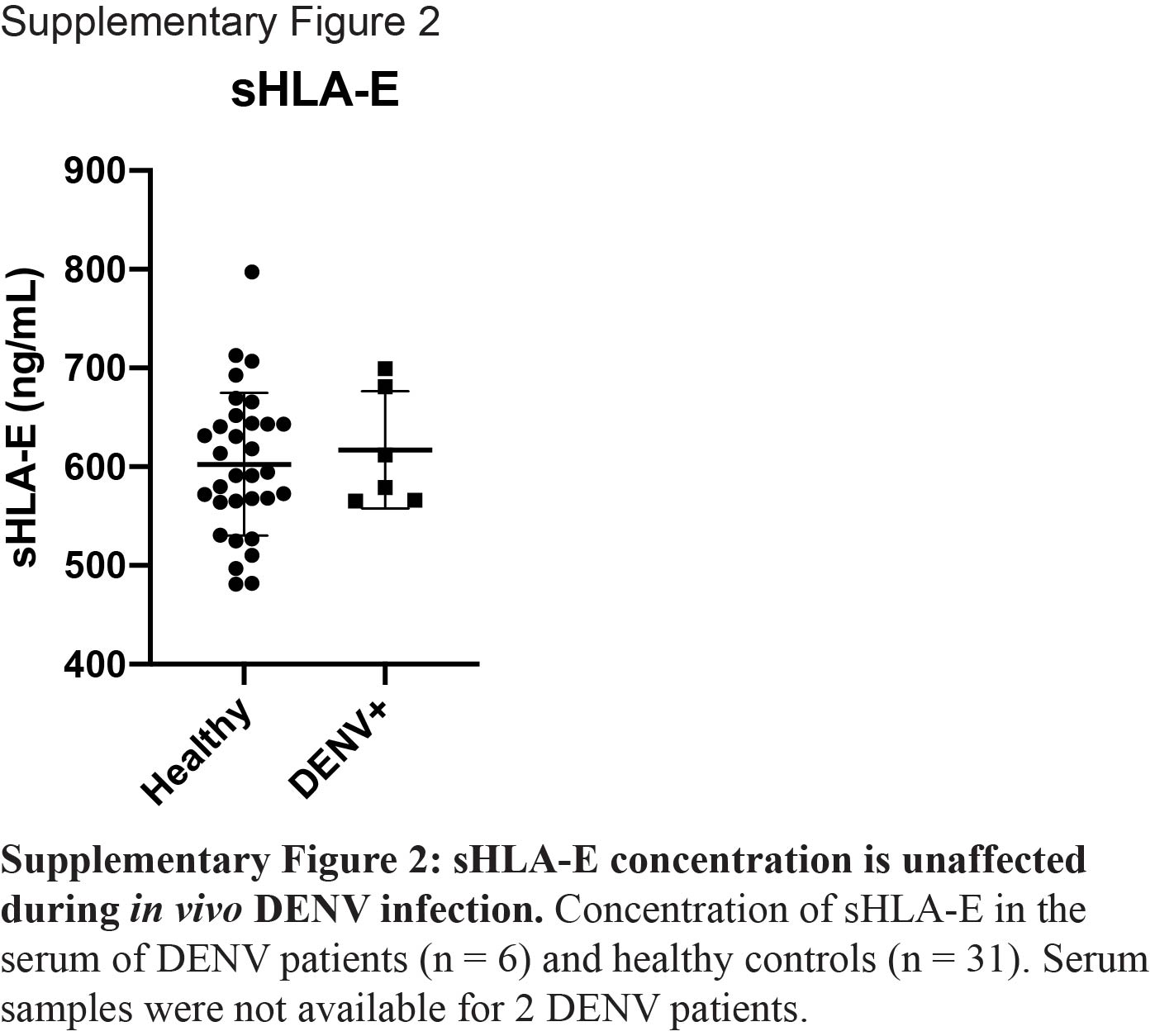

### Supplementary Figure 3

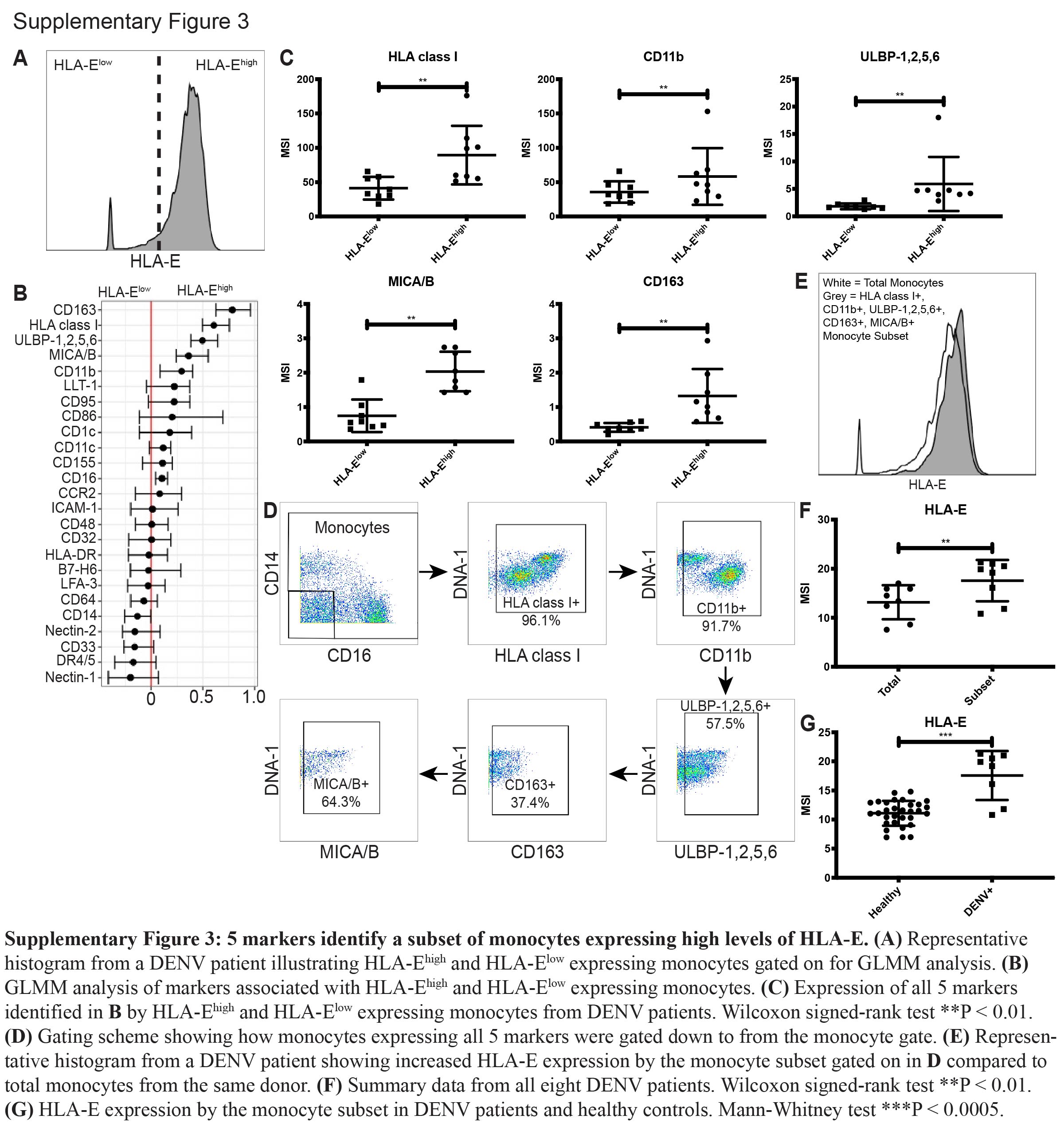

### Supplementary Figure 4

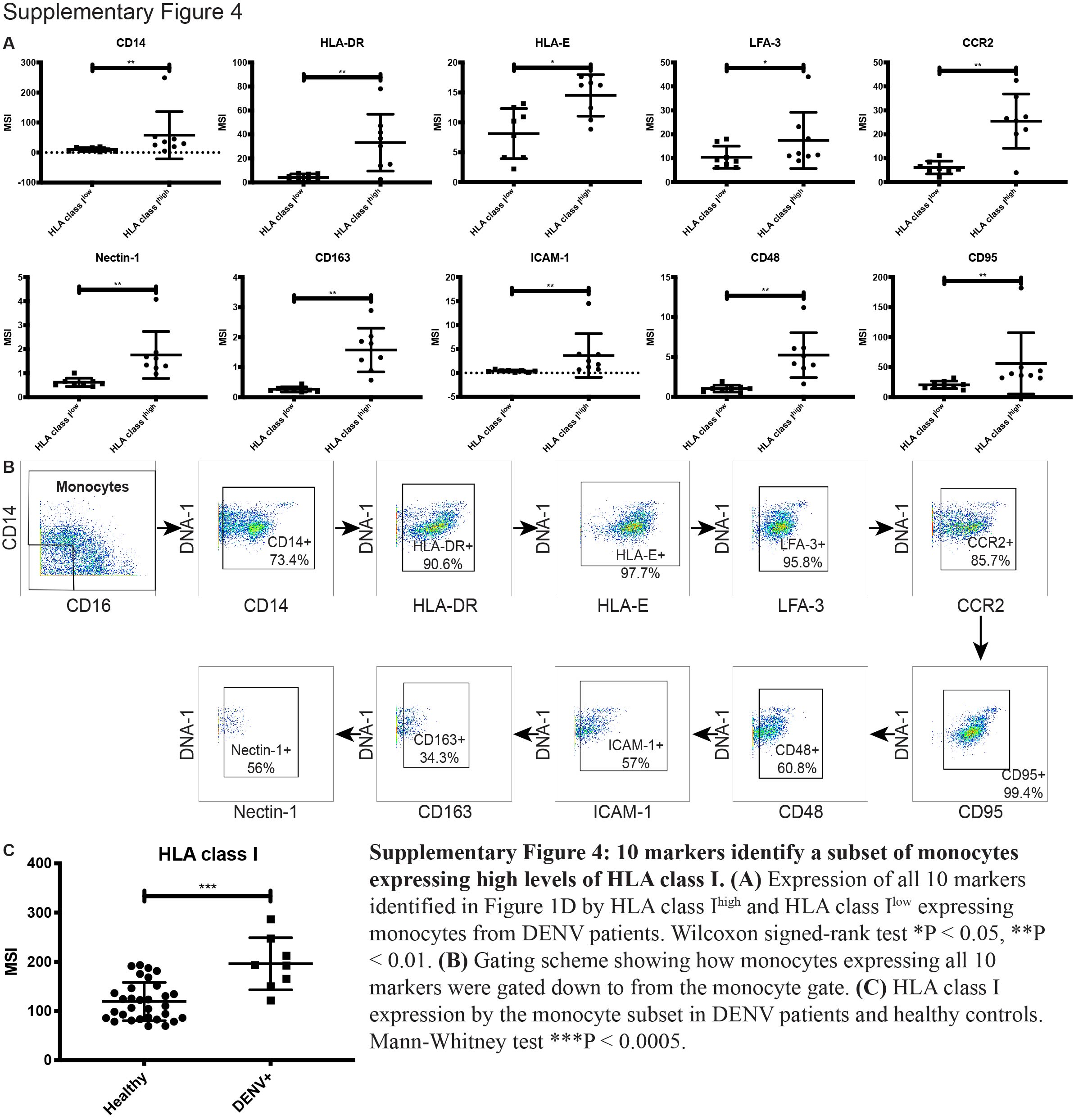

### Supplementary Figure 5

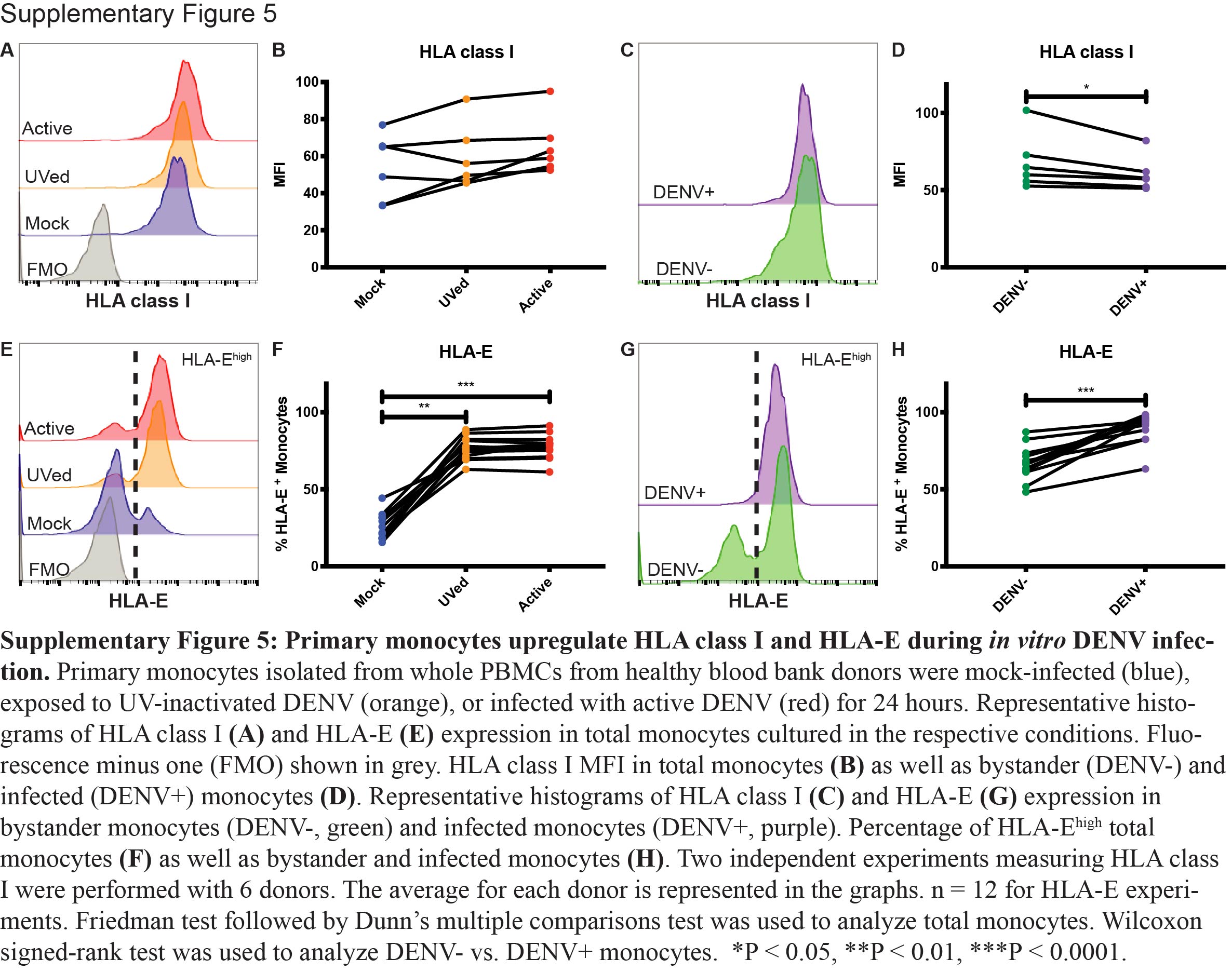
