## Supplementary Table 1 for "HLA upregulation during dengue virus infection suppresses the natural killer cell response"

**Supplementary Table 1:** Summary of dengue study population.

| Characteristic | DENV+ (n = 8) | Healthy (n = 31) |
| --- | --- | --- |
| Age, y, median (range) | 29 (21 - 47) | 31 (16 - 57) |
| Females | 3 | 19 |
| Males | 5 | 12 |
| Days of symptoms, median | 3 | N/A |
